# A molecular phenology scale of fruit development

**DOI:** 10.1101/2022.05.10.491408

**Authors:** Giovanni Battista Tornielli, Marco Sandri, Marianna Fasoli, Alessandra Amato, Mario Pezzotti, Paola Zuccolotto, Sara Zenoni

## Abstract

Fruit growth and development consists of a continuous succession of physical, biochemical, and physiological changes driven by a genetic program that dynamically responds to environmental cues. Establishing recognizable stages over the whole fruit lifetime represents a fundamental requirement for research and fruit crop cultivation. This is especially relevant in perennial crops like the grapevine to scale the development of its fruit across genotypes and growing conditions.

In this work, molecular-based information from several grape berry transcriptomic datasets was exploited to build a molecular phenology scale (MPhS) and to map the ontogenic development of the fruit. The proposed statistical pipeline consisted in an unsupervised learning procedure yielding an innovative combination of semiparametric, smoothing and dimensionality reduction tools. The transcriptomic distance between fruit samples was precisely quantified by means of the MPhS that also enabled to highlight the winding dynamics of the transcriptional program over berry development through the calculation of the rate of variation of MPhS stages by time.

The MPhS allowed the alignment of time-series fruit samples proving to be a step forward in mapping the progression of grape berry development with higher precision compared to classic time- or phenotype-based approaches and inspiring the use of the transcriptional information to scale the developmental progression of any organ in any plant species.

## Introduction

The ontogenic development of fleshy fruits entails an ordered sequence of physical, chemical, physiological, and molecular changes that progressively drive the entire organ towards maturation. These processes are rather conserved among fruits of the same species, but the developmental progression may sway due to both genetic and environmental factors. Moreover, developmental expression patterns are extremely dynamic and, especially under fluctuating environmental conditions (Menzel et al., 2006; Nissanka et al., 2015; Alderman and Stanfill, 2017), rapid changes may happen within short time windows, challenging the setup of meaningful comparisons to study the fruit response to any factor. In annual or perennial fruit crops, the ontogenic development of the fruit is tracked by adopting phenological scales. These are classification tools to describe seasonal and precisely recognized stages of fruit growth and development based on specific descriptors such as visual/physical traits or easy-to-measure compositional parameters (Gillaspy et al., 1993; Labadie et al., 2019). Phenological scales are widely used in models describing known or hypothetical cause-effect relationships between growth stages progression and environmental driving factors (Hess, 1997; Parker et al., 2011; Parker et al., 2020).

In grapevine, the fruit phenophases comprise a large dedicated section of the most adopted phenology scales, namely the modified Eichhorn and Lorenz (E-L) (Coombe, 1995) and the extended BBCH systems (Lorenz et al., 1995), that assign increasing numbers to the main fruit developmental stages from setting to maturity. Grape berry development entails a curve consisting in two phases of growth and lasts up to 150 days (Coombe, 1992). The first phase (also called berry formation or herbaceous/green phase) involves pericarp growth due to rapid cell division and elongation. The second phase (ripening) involves physical and metabolic changes, including softening, skin pigmentation, accumulation of sugars, loss of organic acids, and synthesis of volatile aromas (Conde et al., 2007). The onset of ripening (veraison for viticulturists) occurs after a short lag phase, when the seeds maturation is complete (Coombe, 1992; Fasoli et al., 2018). Growth stages are defined by the assessment of visual/physical traits, such as color, size, and softness. Only for ripening stages and harvest decision, compositional parameters like sugar concentration in the juice are considered. However, the precise definition of developmental stages can be challenging as the fruit traits used for stage description are highly influenced by several factors like genotype, climate, water availability, agronomical practices, and crop load (Sadras and Moran, 2012; Parker et al., 2015; Pastore et al., 2017).

The advent of next-generation sequencing represents an opportunity to exploit the expression kinetic of large sets of genes to stage the fruit development and enrich the available classification systems incorporating information at molecular level (Chuine and Regniere, 2017; Gildor and Smadar, 2018). This approach succeeded to develop a transcriptomic aging clock defining the biological age of organisms such as *C. elegans* to an unprecedented accuracy (Meyer and Schumacher, 2021) and, in plants, was used to reconstruct the transcriptional ontogeny of single organs and correlate the appearance of morphological characteristics with molecularly defined developmental stages (Leiboff and Hake, 2019). Moreover, results obtained on model organisms such as Arabidopsis, or annual crops such as rice, revealed seasonal patterns of gene expression controlled by environmental cues (Nagano et al., 2012; Nagano et al., 2019) and demonstrated that the recent advancement of the methods for gene expression quantification could be exploited to refine phenological stage classification (Kudoh, 2016).

The huge amount of transcriptomic studies generated in recent years over grape berry development revealed that the variation of a portion of the fruit transcriptome is conserved across cultivars and growing conditions (Fasoli et al., 2012; Dal Santo et al., 2013; Robinson et al., 2015; Massonnet et al., 2017; Fasoli et al., 2018; Theine et al., 2021), and thus may be utilized to boost the description of the fruit developmental stages with a molecular dimension.

In this work we used the most informative portion of several grapevine fruit transcriptomic datasets (Massonnet et al., 2017; Fasoli et al., 2018) to build a molecular phenology scale (MPhS). The performance of the scale in precisely mapping the progression of fruit development, over other classical time- or phenotype-based approaches was assessed. The MPhS was used to reinterpret previously published transcriptomic datasets evidencing its potential to trace and compare fruit developmental stages from different genotypes and growing conditions.

## Results

To create a molecular scale of grapevine berry development we relied on the RNA-sequencing dataset consisting of 219 samples published by Fasoli et al. (2018). Samples were collected from fruit set to full maturity from Cabernet Sauvignon (CS) and Pinot noir (PN) vines every 7–10 days across three years (Table S1).

Vintage variability impacted ripening progression since its onset that happened earlier in 2013 and 2014 compared to 2012, and then determined a substantial advance until the grape technological maturity monitored by sugar accumulation (Fig. 1a,b). This behavior was more evident in PN, an early ripening variety respect to CS.

**Fig. 1.**
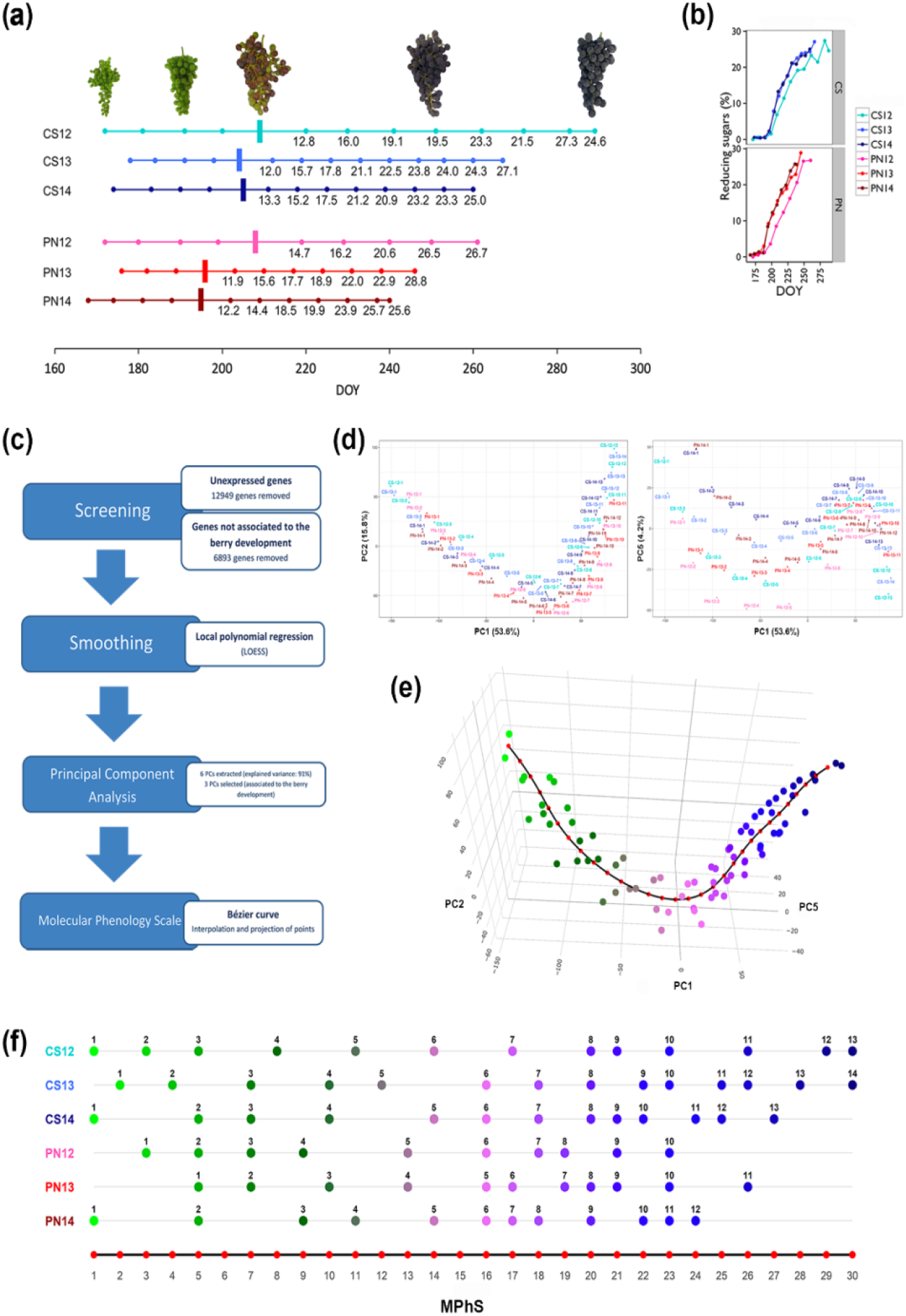
Molecular phenology map creation. (a) Cabernet Sauvignon and Pinot noir time series of berry sample collection. Berries were collected using a randomized block approach to account for intra-vineyard block variability. This resulted in a collection of 219 samples. Brix values are specified for time points post veraison (marked by vertical bar) along the timeline. Veraison: visually defined as the 50%-colored berry in the bunch. DOY, day of the year. Images on top show samples depicting five stages of grape cluster development for Cabernet Sauvignon during season 2012. (b) Accumulation trends of reducing sugars (%) by DOY in Cabernet Sauvignon and Pinot noir samples over three years. Plots were generated using R package ggpplot2 version 2.2.1 (Wickham, 2009). DOY, day of the year. (c) Flow chart of the pipeline. The chart represents the four principal steps (blue rectangles) and related details (white rectangles) (d) Samples distribution according to the three selected PCs. PC1 (53.6%) by PC2 (15.8%) (left) and by PC5 (4.2%) (right). (e) Three-dimension scatterplot of the three selected PCs interpolated by the Bézier curve (black line). Red dots along the Bézier curve define a set of 30 evenly spaced molecular stages. Scattered points correspond to the smoothed samples and changing color highlight berry progression from immature to ripe stages. An interactive plot of the curve is accessible at the link https://bodai.unibs.it/grapevine-gea/mphs/. (f) Projection of the smoothed three-year time series of CS and PN to the closest point among the 30 points identified along the Bézier curve. The sample number is reported above each point. Red dots correspond to the 30 steps of the Molecular Phenology Scale.

Transcriptomic data was analyzed using statistical and data mining tools to define the core set of genes marking the berry development progression (Fig. 1c). The pipeline comprised an initial screening to discard genes exhibiting low and/or noisy expression that reduced the dataset from 29,971 to the whole core set (WCS) of 10,129 genes (Dataset S1), followed by the application of a local polynomial regression that allowed smoothing the gene expression patterns over the time points and averaging the replicates. We then performed Principal Component Analysis (PCA) with the data matrix obtained by column-standardization of the smoothed gene expression. We selected Principal Components (PCs) 1 and 2 that well described the general progression of berry development, and PC5 that improved the discrimination of early stage samples (Fig. 1d), whereas the effect of genotype and vintage (PC3, 4 and 6) was excluded (Fig. S1).The PCs defined a three-dimensional scatter of points, with each point corresponding to an experimental condition (one time point of one cultivar in one year) that were then fitted by one-dimensional space using a Bézier curve. Thirty marks were evenly distributed along the curve to represent steps of what we called Molecular Phenology Scale (MPhS) (Fig. 1e). The points of the 3D scatter were projected onto the MPhS and assigned to the closest among the set of 30 marks, demonstrating that the chronological order of all sampling series was maintained along the MPhS (Fig. 1f), and the core gene set represented a suitable selection for our purpose.

The performances of the MPhS were next explored by projecting the non-smoothed three-year time series of CS and PN on the MPhS. We found that the most part of the samples were correctly ordered according to their chronological collection (Fig. 2a and Table S2). Four sample pairs of CS mapped at the same MPhS stages and in two cases each genotype samples collected at the beginning or at the end of series showed mapping discrepancy with the scale sequence, suggesting a lower resolution of the MPhS at phases with reduced time-related transcriptomic changes. In fact, the samples were not evenly distributed along the MPhS and some intervals (e.g., between MPhS stage 7 and 12) were poorly represented. The veraison phase was pinpointed in correspondence to the MPhS window 13-15 for both genotypes, whereas PN always reached maturity at earlier MPhS stage compared to CS.

**Fig. 2.**
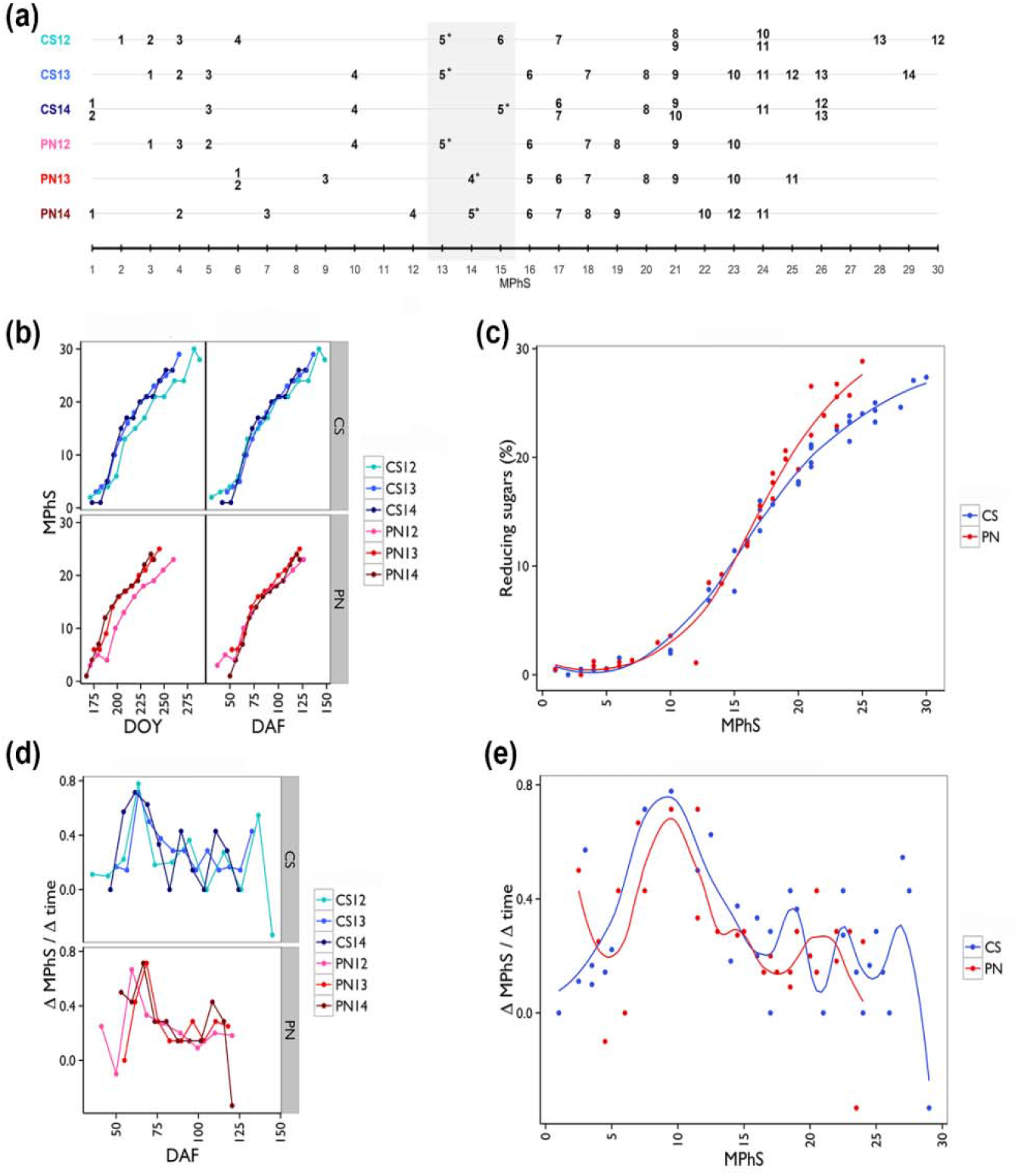
Relationship between Molecular Phenology Scale and time during fruit development. (a) Projection of the non-smoothed three-year time series of CS and PN on the MPhS. Samples collected at version phase are marked with an asterixis. The light-grey rectangle highlights the MPhS stages corresponding to the version transition. (b) Trends of MPhS stages by day of the year (DOY) (left) and day after flowering (DAF) (right) in CS and PN over the three years. Plots were generated using R package ggpplot2 version 2.2.1 (Wickham, 2009). (c) Trend of percentage of reducing sugars accumulation by MPhS in CS and PN. Smoothed conditional means function of the R package ggpplot2 version 2.2.1 (Wickham, 2009) was used to represent the average of the three years per genotype. (d) Plot of the ΔMPhS/Δtime over DAF. ΔMPhS represents the difference in MPhS between subsequent stages. Δtime represents the difference in days between subsequent stages. Plots were generated using R package ggpplot2 version 2.2.1 (Wickham, 2009). (e) Plot of the ΔMPhS/Δtime over the MPhS. Smoothed conditional means function of the R package ggpplot2 version 2.2.1 (Wickham, 2009) was used to represent the average of the three years per genotype.

Plotting the MPhS by the day of the year (DOY) confirmed differences among years and evidenced a transcriptional progression delayed in 2012 in both varieties, whereas the alignment on the phenological flowering phase resulted in nearly overlapping curves unraveling common kinetics, with the very first phase characterized by slow transcriptional changes followed by a rapid advancement clearly anticipating the veraison stage (Fig. 2b). Because similar dynamics were observed when the MPhS series were plotted on the heat summation, we speculate that temperature is less effective than time in driving fruit transcriptomic progression (Fig. S2).

When exploring the relation between MPhS stage and sugar content of the PN and CS samples, we observed a strict non-linear relation that clearly varied in the two genotypes, with PN reaching technological maturity at earlier MPhS stages than CS (Fig. 2c). On the other hand, an elevated variability, beyond the clear genotype effect, was observed between MPhS and Berry weight, suggesting that the two variables are poorly related (Fig. S3).

The transcriptomic scale dynamics highlighted by the ratio ΔMPhS/Δtime (an approximation of the derivative of the MPhS curve over time) showed clear fluctuations irrespective of the DOY (Fig. S4) and more aligned by genotype when plotted over the day after flowering (DAF), evidencing a major peak likely associated with the onset of ripening, followed by some minor peaks (Fig. 2d).

This dynamic was better highlighted by plotting the ΔMPhS/Δtime averaged by year over the MPhS itself, revealing that some MPhS stages are rapidly passed through (Fig. 2e). A main burst of speed was reached at stages 9-10 in both genotypes, followed by minor accelerations at stages 17-18, 22-23 and 27-28 in CS and around stage 22 in PN.

The performance of the MPhS was then tested on previously published berry transcriptomes by mapping the relative samples onto the MPhS using the core set of 10,129 genes selected for the scale definition.

The RNA-seq dataset from Massonnet et al. (2017) provided information in ten genotypes across four berry phenological stages (BBCH scale) of the same season. When scaled onto the MPhS, the two early stages, Pea Size and Touch, mapped nearby between stage 6 and 11 (Fig. 3a). Later stages appeared less aligned across genotypes, spanning from the white-skin variety Passerina that mapped at MPhS stage 20 at Harvest, to the red-skin Barbera, Negroamaro and Refosco, exhibiting maturity at 25.

**Fig. 3.**
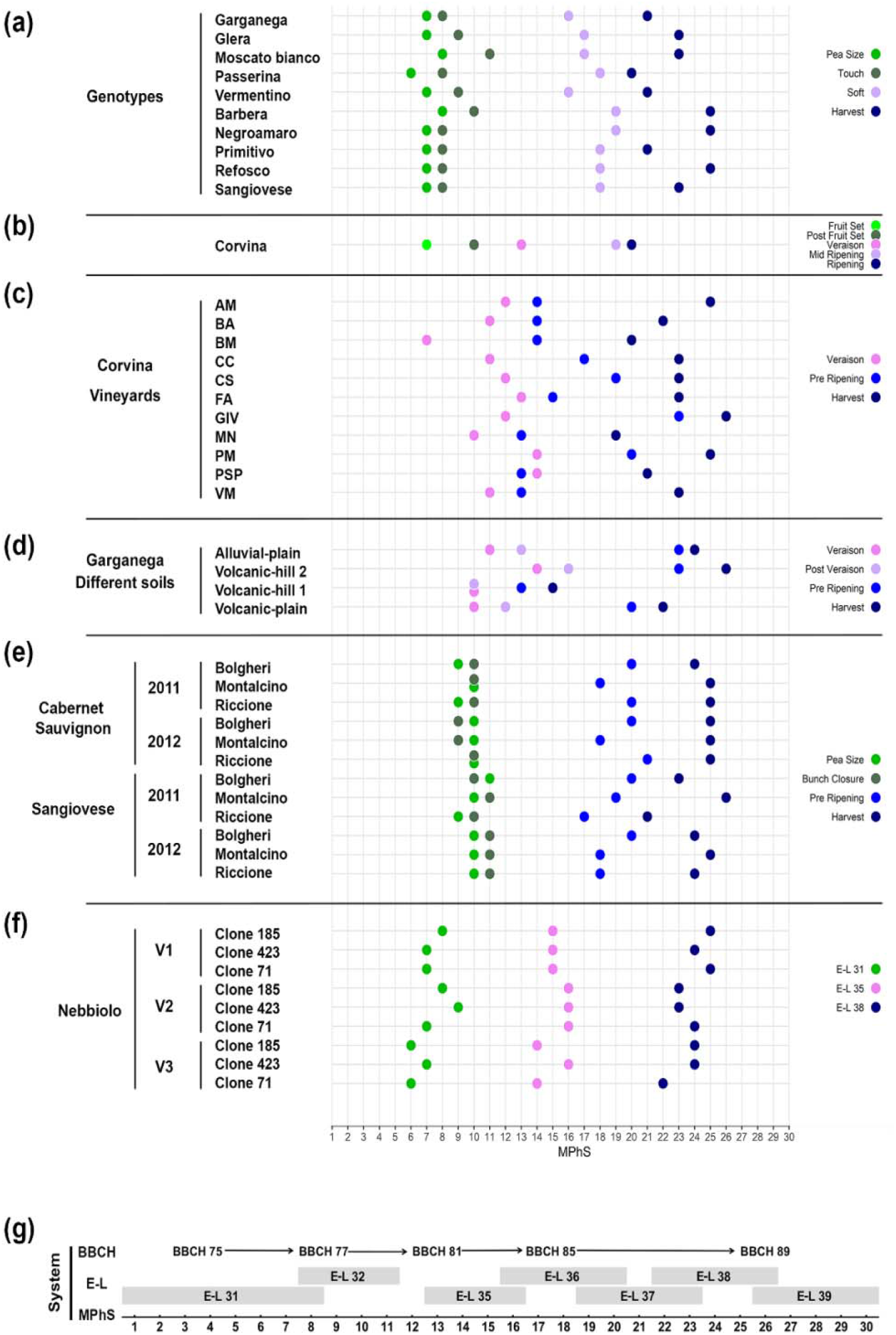
Scaling of previously published berry transcriptomic samples onto the MPhS. (a) Projection of RNA-seq transcriptomic berry samples of ten varieties (Massonnet et al., 2017) onto the MPhS. The BBCH scale was followed to collect berry samples at specific phenological stages: Pea Size (BBCH 75), Touch (BBCH 77), Soft (BBCH 85) and Harvest (BBCH 89) during the same season. (b) Projection of microarray transcriptomic berry samples of cv Corvina (Fasoli et al., 2012) onto the MPhS. Samples collection was performed at phenological stages defined by the E-L scale: Fruit Set (E-L 29), Post Fruit Set (E-L 32), Veraison (E-L 35), Mid Ripening (E-L 36) and Ripening (E-L 38). (c) Projection of microarray transcriptomic berry samples of cv Corvina collected at three ripening stages (Veraison, Pre Ripening and Harvest) from eleven cultivation sites (Dal Santo et al., 2013), onto the MPhS. (d) Projection of microarray transcriptomic berry samples of cv Garganega collected at four ripening stages (Veraison, Post Veraison, Pre Ripening and Harvest) from four cultivation sites (Dal Santo et al., 2016), onto the MPhS. (e) Projection of microarray transcriptomic berry samples of cv Sangiovese and cv Cabernet Sauvignon compared across three growing sites and over two years (Dal Santo et al., 2018), onto the MPhS. Berries were collected at four developmental stages defined by the BBCH phenological scale: Pea Size (BBCH 75), Bunch Closure (BBCH 79), Pre Ripening (BBCH 83) and Harvest (BBCH 89). (f) Projection of RNA-seq transcriptomic berry samples of three clones of cv Nebbiolo, compared across three different sites (Pagliarani et al., 2019), onto the MPhS. Samples were collected at three developmental stages defined by the E-L phenological scale: Pea Size (E-L 31), Veraison (E-L 35), and Ripening (E-L 38). We represented seven main stages in the legend: stage 1 (Fruit Set and E-L 29; light green), stage 2 (Pea Size and E-L 31; green), stage 3 (Touch, Post Fruit Set, Bunch Closure and E-L 32; olive green), stage 4 (Veraison and E-L 35; pink), stage 5 (Soft, Mid Ripening, Post Veraison and E-L 36; plum), stage 6 (Pre Ripening and E-L 37; blue) and stage 7 (Harvest, Ripening and E-L 38, midnight blue). (g) Alignment between MPhS and two phenotype-based phenological scales (BBCH and the modified E-L) during berry development.

We also scaled samples from microarray-based transcriptomic datasets among which the five phenological stages (E-L scale) of the cultivar Corvina (Fasoli et al., 2012). Albeit obtained by a different transcriptomic technology, samples mapped neatly onto the MPhS, with green berry samples projected at stage 7 and 10, Veraison at stage 13, whereas Mid Ripening and Ripening at stages 19 and 20, respectively (Fig. 3b).

A benefit of the molecular scale consists in recalibrating studies that organized sampling on a time-based approach rather than following berry phenology. This was the case of two microarray-based transcriptomic datasets in which berries of the cultivar Corvina were collected at three ripening times from eleven sites (Dal Santo et al., 2013), whereas for the cultivar Garganega they were collected at four ripening times from four sites (Dal Santo et al., 2016). As expected, datasets projection onto the MPhS revealed a fair misalignment of time points collected from different sites (Fig. 3c,d). The greatest differences were observed for Corvina samples at the second stage ranging from MPhS stage 13 to 23, whereas Garganega berries mapped between MPhS stage 15 and 26 at harvest.

Mapping samples from the work of Dal Santo and co-authors (2018), exploring the genotype by environment interaction (GxE), revealed that berries of the cultivar Sangiovese reached maturity at different MPhS stages by cultivation site and year whereas Cabernet Sauvignon samples appeared much more aligned (Fig. 3e) affirming the marked transcriptomic plasticity of Sangiovese. The strength of the MPhS mapping approach to GxE was also assessed when the genotype component represented different clones, like in the work of Pagliarani et al. (2019) entailing RNA-seq berry samples of three clones of the cultivar Nebbiolo, grown in three sites, and collected at three developmental stages. Although samples arranged along the MPhS by E-L phenological classification (Fig. 3f), minor MPhS shifts reflecting the intrinsic transcriptomic plasticity of each clone interacting with the growing site were yet appreciable.

The available sampling metadata was then exploited to attempt an alignment between our MPhS and the classical phenotype-based phenological scales (i.e., the modified E-L and BBCH systems) during berry development (Fig. 3g), showing the greater classification detail provided by the MPhS compared to the traditional scales.

Starting from the core set of genes used to create the MPhS, we next focused on identifying the smallest number of genes necessary to efficiently map berry transcriptomic samples onto the MPhS.

Pools of 20, 10, 5 and 2 positive and negative loadings of each of the three PCA components (corresponding to 120, 60, 30 and 12 genes, respectively) were selected based on their absolute correlation value (p-corr) and expression profile. Hierarchical cluster analysis of the top-100 loadings throughout berry development in PN and CS during the three vintages revealed 20 main clusters of gene expression (Fig. S5). As genes belonging to the same metabolic/developmental process likely co-express, we picked genes with high p-corr value from each cluster to maximize the information potential thus avoiding redundancy (Dataset S2). Calculating the average shift between the projections of the sample series defined by the reduced core sets (RCS) and the WCS allowed evaluating any difference in performance (Fig. 4). This analysis highlighted that the progressive reduction of the core gene set from 120 to 12 did not substantially impacted the samples order along the MPhS and only slightly worsened its scaling performance.

**Fig. 4.**
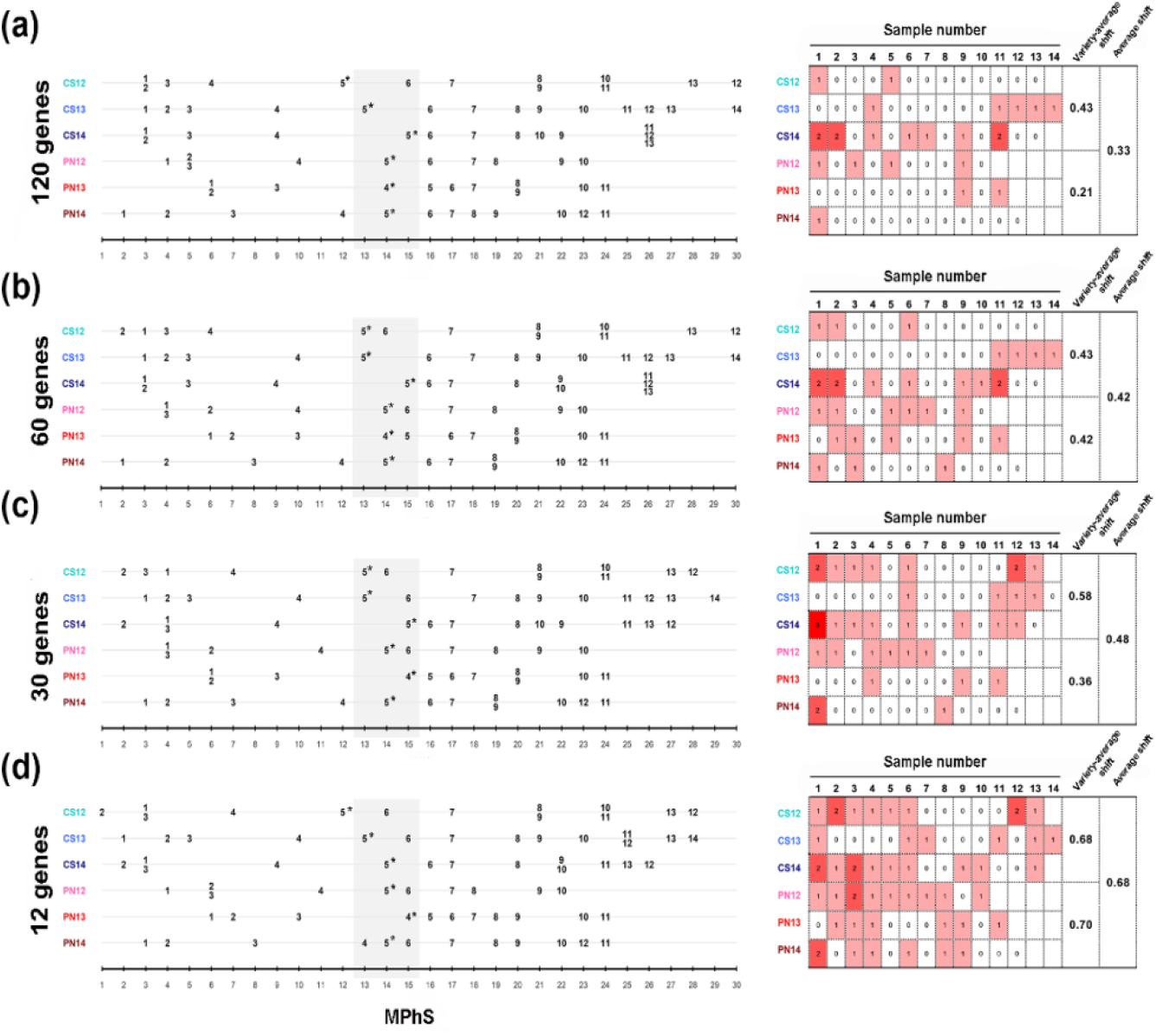
Performance of MPhS based on a reduced core set of genes. Projections of the sample series defined by the progressively reduced core set of genes (a) 120 genes, (b) 60 genes, (c) 30 genes and (d) 12 genes (Dataset S2) onto the MPhS (left) and representation of the average shifts between the distribution of samples defined by reduced core sets versus when defined by the entire core set of genes (10,129) (right). For each sample the shift between the MPhS stage defined by the reduced gene core set and the MPhS stage defined by the entire gene core set was calculated. The overall and by variety average shift are shown per reduced core set of genes.

## Discussion

Assembling meaningful comparisons between fruits characterized by different developmental rates or seasonal developmental shifts is a major limitation in studying the biological mechanisms underpinning the seamless developmental progression that leads the fruit to maturation. The opportunity of refining the existing development classification systems by integrating phenotype with molecular-based information has been explored in model organisms or annual crops (Kudoh, 2016; Nagano et al., 2019; Meyer and Schumacher, 2021). The transcriptomic analysis of immature maize tassels and sorghum panicles throughout development enabled reconstructing their transcriptional ontogeny and correlating the emergence of species-specific morphological characteristics of these organs to developmental stages defined at molecular level (Leiboff and Hake, 2019). Despite omics technologies have been widely applied to unveil and describe in detail the ongoing developmental changes in fleshy fruits from many species (Janssen et al., 2008; Sanchez-Sevilla et al., 2017; Fasoli et al., 2018; Shinozaki et al., 2018), the molecular information was rarely exploited to attempt at scaling the fruit ontogeny. In grapevine, Dai et al. (2013) demonstrated that metabolite profiling can be used in PCAs to define development trajectories for berries, whereas Wang et al. (2014) showed that the expression trend of a small number of genes involved in grapevine flower and berry development was associated to some specific phenological stages. In this study, we selected about ten thousand genes from several transcriptomic datasets for their consistent expression throughout berry development regardless genotype and season (Massonnet et al., 2017; Fasoli et al., 2018) and built a molecular phenology scale to map the ontogenetic development of the fruit with high precision and to align berry development of different grapes. The proposed statistical pipeline consisted in an unsupervised learning procedure yielding an innovative combination of semiparametric, smoothing and dimensionality reduction tools. Time information was exploited by only considering timepoints succession disregarding their distance so the MPhS units are not time but ideal steps of the berry development, which can take longer or shorter, providing the flexibility of accounting for a multiplicity of factors. Additionally, it is worth mentioning that the Principal Components were exceptionally able to summarize different characteristics of the data (berry variety, vintage, stage) allowing to precisely select genes involved in berry development and rule out those impacted by genotype and vintage.

When projecting onto the MPhS the sample series used for its own constructions, the distribution was largely by the time of collection during the season, with very few overlaps or inversions. These cases were mainly concentrated at the initial and final stages of the scale suggesting that berry transcriptome evolution right after fruit set and near maturity is slower than during the central time frame, therefore samples collected at two consecutive weeks could exhibit very similar transcriptomes. Indeed, the uneven distribution of samples along the scale mirrored the winding dynamics of berry transcriptome over development revealing that some MPhS stages are more rapidly crossed by the developing fruit. For both varieties the highest rate of variation of the MPhS stage by time (represented by a major peak) was recorded well prior the assessed veraison stage, confirming that the ripening transcriptional program is established one-to-two weeks before berry phenotypic changes can be visually appreciated (Castellarin et al., 2016; Fasoli et al., 2018; Hernández-Montes et al., 2021). Following the onset of ripening, minor peaks were present towards maturity suggesting that the ripening progression is discontinuous at transcriptional level. We exclude these fluctuations to be caused by interactions with the environment as the statistical pipeline was set to screen out the vintage component and any erratic variation of gene expression.

Applications of the MPhS are aligning samples to highlight shifts of fruit development driven by factors like genotype and environment as well as precisely quantifying their transcriptomic distance. This could represent a start to pinpoint the specific contribution of any variance in fruit development and the cultivation environment to the overall plasticity. Once fruit samples align by MPhS stage, comparing differentially expressed genes could unveil responses uniquely related to the growing conditions. Projecting RNA-seq and microarray transcriptomic samples (Fasoli et al., 2012; Dal Santo et al., 2013; Dal Santo et al., 2016; Massonnet et al., 2017; Dal Santo et al., 2018; Pagliarani et al., 2019) onto the MPhS indeed highlighted phenological differences and provided the basis to shed the light into instances when samples originally assigned to similar phenological stage mapped relatively apart along the MPhS. Nevertheless, the MPhS ability to discriminate samples collected at a considerable time distance was not fully confirmed in certain dataset/time series. This may be related to specific raw data processing and procedures adopted in each study, prompting the need of future investigation and development of broader normalization protocols to improve the performance of the MPhS.

Regarding the classification of ripening stage, the molecular scale represents an advance over the traditional analytical methods (i.e., total soluble solids or reducing sugars percentage in the grape juice) that, although rapid, do not always represent a reliable indicator of the berry physiological ripening stage. In fact, the great influence of genetics, climate, and agronomic factors on sugar accumulation dynamics in the fruit can lead to a partial uncoupling from other ripening technological parameters, like organic acids and skin pigment content (Sadras and Moran, 2012; Parker et al., 2015; Pastore et al., 2017). We found a close relation between MPhS stage and berry sugar level at initial stages of maturation in both cultivars, that diverged at late maturation likely reflecting the different length of ripening characterizing the two genotypes and indicating that CS berries reached the target level of soluble solids at an advanced MPhS stage compared to PN.

Based upon the metadata available for each dataset, we attempted to approximate the 1-30 MPhS stages to those specified in the classical grapevine fruit phenology scales and we thus attested that the MPhS permits greater definition of the phenophases respect to the classical scales (Coombe, 1995; Lorenz et al., 1995) that cover longer stretches along berry development.

Finally, we showed that the number of expression signals necessary for mapping samples on the MPhS can be drastically reduced to a few dozen without substantial loss of precision, creating the opportunity of defining the molecular phenology stage of a fruit sample just performing targeted gene expression analysis.

The proposed scale paves the way for the development of tools that aspire at predicting the phenological stage of the fruit in various climate conditions, such as models that can account for temperature and other environment clues. The quality of these tools will benefit from the combination of various modeling techniques (molecular, metabolite, physical, visual levels), providing that great coordination and knowledge-transfer between modelers, biologists and growers will be established.

The proposed pipeline could be potentially extended and successfully applied to any other fruit species, provided they have some basic requirements: (i) a relatively frequent sampling covering the time-series transcriptomic changes during fruit development with high detail; (ii) the availability of expression data from diverse growing conditions and genotypes allowing the phenology scale to be representative of the general development of the fruit of the species; (iii) a reliable and well annotated reference genome to compute the expression data. The existence of these conditions ensured the successful implementation of our MPhS for the grapevine berry phenological classification, that we foresee will help coping with challenges such as those raised by climate change, allowing the precise mapping of the berry developmental progression.

## Materials and Methods

### Transcriptomic datasets

The expression data used for the creation of MPhS was retrieved from Fasoli et al. (2018) (data accession number GSE98923), whereas the berry transcriptomes projected onto the MPhS referred to: Massonnet et al. (2017) (data accession number GSE62744 and GSE62745); Fasoli et al. (2012) (data accession number GSE36128), Dal Santo et al. (2013) (data accession number GSE41633); Dal Santo et al. (2016) (data accession number GSE75565); Dal Santo et al. (2018) (data accession number GSE97578); Pagliarani et al. (2019) (data accession number GSE116238).

### Data mining process

We used the data described in Fasoli et al. (2018) (*dataset A*) and set the following variables:

- Cultivar [Cabernet Sauvignon, Pinot Noir]
- Vintage [2012, 2013, 2014]
- Timepoint [from 1 to 10-14 (different combinations Cultivar x Vintage have different final Timepoints)]

*C*x*V* denotes the six possible combinations Cultivar x Vintage and the term *experimental condition* represents each of the 73 possible combinations Cultivar x Vintage x Timepoint.

Genetic Variables:

- Gene Expression [RNAseq platform]

*x_jri_* denotes the expression of gene *i* (*i*=1, 2, …, 29,971) in the *j*th experimental condition (*j*=1, 2, …, 73) for the *r*th replicate (*r*=1, 2, 3); the subscript *h* represents the combination *C*x*V* (*h*=1, 2, …, 6). *J_h_* denotes the set of experimental conditions corresponding to *C*x*V* = *h*.

In addition, *m_ji_* indicates the average expression (over the three replicates) of gene *i* in the *j*th experimental condition

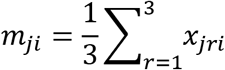

and *m_hi_* the average expression (over the replicated and the timepoints) of gene *i* in the *h*th *C*x*V* combination

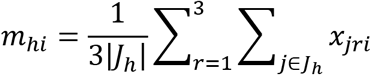

where |·| denotes cardinality of a set.

Th expression profiles of each gene were displayed using a three-panel graphical representation (Fig. S6), with one graph each vintage.

The data mining process comprises four steps (Fig. S7):

### Step 1: Screening

We removed 19,842 genes from the 29,971 in the grapevine transcriptome exhibiting uninteresting profiles (i.e., no expression in some experimental conditions or expression not associated to berry development; Fig. S8) based on the criteria summarized in Table S3. To improve the selection, this screening step exploited the information of an additional dataset (*dataset B*), composed of 10 cultivars, observed for 4 timepoints during fruit development in a single vintage (Massonnet et al., 2017). The resulting dataset comprised 10,129 genes (denoted as *F*) deserving further statistical analysis.

### Step 2: Smoothing

Smoothing was applied to the data matrix (219 × 10,129) containing the expressions *x_jri_* with *i* ∈ *F*. For each gene *i* and for each *C*x*V* combination *h*, we estimated smoothed values 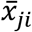 by means of a polynomial local regression method called LOESS, introduced by Cleveland (1979) and Cleveland and Devlin (1988). LOESS fits a low-degree polynomial to a subset of the data in the neighborhood of each observation of the dataset. In other words, simple parametric models are fitted to localized subsets of the data, aiming at obtaining a smooth curve. The polynomial parameters are estimated by a weighted least squares method, where a higher weight is given to points closer to the observation whose outcome is being calculated. The fraction of the total number of data points that were used in each local fit is determined by the smoothing parameter, usually denoted by α.

In the context of this application, for a gene *i*, the single observation was the expression of replicate *r* of a given cultivar, in a given vintage, at a given timepoint. The neighborhood was composed of all the expressions of gene *i*, for the same cultivar and in the same vintage, in the nearby timepoints, with higher weights to the closest ones. We used polynomials of degree 2 and set α = 0.75. The outcome 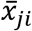 for the experimental condition *j* ∈ *J_h_* was then calculated by evaluating the local polynomial at the timepoint characterizing the experimental condition.

For each gene, a three-panel graphical representation could be visualized, analogous to that presented in Fig. S6, where, in each panel, the line shows the pattern of the smoothed gene expression 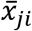 (Fig. S9).

### Step 3: Principal Component Analysis

This step was applied to the data matrix (73 × 10,129) containing the smoothed expressions 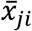 with ∈ *F*. We performed PCA (Jolliffe, 1986) on the standardized data matrix and extracted six PCs accounting for a 91% explained variance. The interpretation of the PCs was based on the graphical representations in Fig. S10, where the pattern of each PC was plotted against the timepoints, with lines corresponding to the different *C*x*V* combinations.

### Step 4: Molecular Phenology Scale definition

This step concerned the data matrix (73 × 3) containing the values of PC1, PC2 and PC5 for each experimental condition, which can be represented as a scatter of points in a three-dimensional Euclidean space, with the implicit additional information of a timeline expressed by the timepoints and the calendar day. The scatter of points was interpolated by means of a Bézier curve (Rabut, 2002), which nowadays is employed in several applications, especially in the field of computer graphics. A Bézier curve aims at fitting points with a smooth curve completely contained within the convex hull of a set of *k* control points (*k* is called the curve’s order): the first and last control points are the end points of the curve, whereas the curve does not pass through the intermediate ones (if any), which define orientation and shape.

In the context of our application, we set *k* = 5 and obtained the curve in Fig. S11. An interactive plot of the curve is accessible at the link https://bodai.unibs.it/grapevine-gea/mphs/.

We projected the scatter on the Bézier curve by assigning each point of the scatter to the closest point among a set of 30 evenly spaced points identified along the Bézier curve. We represented the curve as a linear graduation, called Molecular Phenology Scale. Each experimental condition was assigned to one of the 30 marks and the Molecular Phenology Scale was represented for each *C*x*V* combination *h*, thus showing on the map only the experimental conditions *j* ∈ *J_h_*.

### Projection of transcriptomic data on the MPhS

In this section we describe the procedure to project observations coming from different case studies onto the Molecular Phenology Scale. We used the matrix *A* of the eigenvectors of the three selected Principal Components used to build the Molecular Phenology Scale.

Let *Z_obs_* = {*z_ji_*} be a |*J_obs_*| x |*I_obs_*| data matrix containing the observed expression levels of a set *I_obs_* of genes for a set *J_obs_* of experimental conditions, where *j* = 1, 2, ⋯, |*J_obs_*| and *i* = 1,2, ⋯, |*I_obs_*|. The matrix *Z_obs_* was column standardized. We consider two cases:

1. *F* ⊆ *I_obs_* (recall that *F* is the set of 10,129 selected genes after the screening step). In this case we obtained the matrix 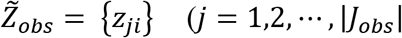 and *i* = 1,2, ⋯, 10129) by removing from *Z_obs_* the columns not belonging to *F*. Using 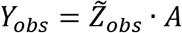 we obtained the estimates of the three Principal Component values for the observed values of the |*J_obs_*| experimental conditions;
2. *I_obs_* ⊂ *F*. This case typically occurs when a limited number of genes is observed, for example for cost reasons. When this is the case, it is recommended to observe genes highly correlated to the Principal Components, that should help obtaining good estimates in spite of the scarcity of data. Starting from the matrix *Z_obs_* containing the observations of a limited number of genes |*I_obs_*|, we obtained the estimates of the Principal Component values for the observed values of the |*J_obs_*| experimental conditions as follows:

- we computed the average expressions of the |*I_obs_*| genes positively and negatively correlated to each Principal Component;
- for Principal Component *q*, we imputed the missing data (non-observed expressions for genes in *F*) with the respective positive or negative average, according to the sign of the corresponding eigenvector coefficient;
- let this imputed |*J_obs_* | × 10129 matrix be denoted with *Z_obs,q_*, we obtained the estimates of Principal Component *q* as *Y_obs,q_* = *Z_obs,q_* · *A_q_*, where *A_q_* was the *q*-th column of *A*.

Once obtained the estimates of the three Principal Component values, we plotted the estimated three-dimensional scatter. This cloud of points was then projected onto the Bézier curve estimated in step 4 by assigning each point to its closest marker among the set of 30 evenly spaced points previously identified along the curve.

### Selection of transcripts for the reduced core set-based scale

We investigated four scenarios in case 2 (*I_obs_* ⊂ *F*), characterized by different subsets of genes with |I_obs |=120, 60, 30, 12, respectively. For the selection of 120, 60, 30, 12 genes, the expression profile of the top-100 loadings characterized by the highest and lowest correlation values with the three dimensions of the PCA were hierarchically clustered based on average Pearson’s distance metric (Tmev 4.3). We selected genes from each of the 20 different clusters. Genes with the highest p-corr value within a cluster were preferred. For the PC5 the selection was based only on the p-core values given their general low levels.

## Acknowledgments

This work is based upon work from COST Action CA17111 INTEGRAPE, supported by COST (European Cooperation in Science and Technology).

## Author contributions

GBT and SZ conceived the research; GBT, MS, PZ and SZ designed the experiments; MS, PZ and MF performed experiments; MS, PZ, MF, GBT and SZ analysed data; GBT, MS, PZ, SZ, MF interpreted data; MP and AA helped drafting the manuscript; GBT, SZ and MF wrote the manuscript; GBT and MS contributed equally.

## Parsed Citations

Alderman PD, Stanfill B (2017) Quantifying model-structure- and parameter-driven uncertainties in spring wheat phenology prediction with Bayesian analysis. European Journal of Agronomy 88: 1–9 Google Scholar: Author Only Title Only Author and Title

Castellarin SD, Gambetta GA, Wada H, Krasnow MN, Cramer GR, Peterlunger E, Shackel KA, Matthews MA (2016) Characterization of major ripening events during softening in grape: turgor, sugar accumulation, abscisic acid metabolism, colour development, and their relationship with growth. J Exp Bot 67: 709–722 Google Scholar: Author Only Title Only Author and Title

Chuine I, Regniere J (2017) Process-Based Models of Phenology for Plants and Animals. Annual Review of Ecology, Evolution, and Systematics, Vol 48 48: 159–182 Google Scholar: Author Only Title Only Author and Title

Cleveland DW (1979) Robust locally weighted regression and smoothing scatterplots. Journal of the American Statistical Association 74 Google Scholar: Author Only Title Only Author and Title

Cleveland DW (1988) Locally weighted regression: an approach to regression analysis by local fitting. Journal of the American Statistical Association 83 Google Scholar: Author Only Title Only Author and Title

Conde C, Silva P, Fontes N, Dias ACP, Tavares RM, Sousa MJ, Agasse A, Delrot S, Gerós H (2007) Biochemical changes throughout grape berry development and fruit and wine qua. Food 1: 22 Google Scholar: Author Only Title Only Author and Title

Coombe BG (1992) Research on Development and Ripening of the Grape Berry. American Journal of Enology and Viticulture 43: 101–110 Google Scholar: Author Only Title Only Author and Title

Coombe BG (1995) Adoption of a system for identifying grapevine growth stages. Australian Journal of Grape and Wine Research 1: 10 Google Scholar: Author Only Title Only Author and Title

Dai ZW, Leon C, Feil R, Lunn JE, Delrot S, Gomes E (2013) Metabolic profiling reveals coordinated switches in primary carbohydrate metabolism in grape berry (Vitis vinifera L.), a non-climacteric fleshy fruit. J Exp Bot 64: 1345–1355 Google Scholar: Author Only Title Only Author and Title

Dal Santo S, Fasoli M, Negri S, D’Inca E, Vicenzi N, Guzzo F, Tornielli GB, Pezzotti M, Zenoni S (2016) Plasticity of the Berry Ripening Program in a White Grape Variety. Front Plant Sci 7: 970 Google Scholar: Author Only Title Only Author and Title

Dal Santo S, Tornielli GB, Zenoni S, Fasoli M, Farina L, Anesi A, Guzzo F, Delledonne M, Pezzotti M (2013) The plasticity of the grapevine berry transcriptome. Genome Biol 14: r54 Google Scholar: Author Only Title Only Author and Title

Dal Santo S, Zenoni S, Sandri M, De Lorenzis G, Magris G, De Paoli E, Di Gaspero G, Del Fabbro C, Morgante M, Brancadoro L, Grossi D, Fasoli M, Zuccolotto P, Tornielli GB, Pezzotti M (2018) Grapevine field experiments reveal the contribution of genotype, the influence of environment and the effect of their interaction (GxE) on the berry transcriptome. Plant J 93: 1143–1159 Google Scholar: Author Only Title Only Author and Title

Fasoli M, Dal Santo S, Zenoni S, Tornielli GB, Farina L, Zamboni A, Porceddu A, Venturini L, Bicego M, Murino V, Ferrarini A, Delledonne M, Pezzotti M (2012) The grapevine expression atlas reveals a deep transcriptome shift driving the entire plant into a maturation program. Plant Cell 24: 3489–3505 Google Scholar: Author Only Title Only Author and Title

Fasoli M, Richter CL, Zenoni S, Bertini E, Vitulo N, Dal Santo S, Dokoozlian N, Pezzotti M, Tornielli GB (2018) Timing and Order of the Molecular Events Marking the Onset of Berry Ripening in Grapevine. Plant Physiol 178: 1187–1206 Google Scholar: Author Only Title Only Author and Title

Gildor T, Smadar BD (2018) Comparative Studies of Gene Expression Kinetics: Methodologies and Insights on Development and Evolution. Front Genet 9: 339 Google Scholar: Author Only Title Only Author and Title

Gillaspy G, Ben-David H, Gruissem W (1993) Fruits: A Developmental Perspective. Plant Cell 5: 1439–1451 Google Scholar: Author Only Title Only Author and Title

Hernández-Montes E, Zhang Y, Chang B-M, Shcherbatyuk N, Keller M (2021) Soft, Sweet, and Colorful: Stratified Sampling Reveals Sequence of Events at the Onset of Grape Ripening. American Journal of Enology and Viticulture 72: 137–151 Google Scholar: Author Only Title Only Author and Title

Hess M, Barralis, G., Bleiholder, H., Buhr, L., Eggers, TH., Hack, H., Stauss, R. (1997) Use of the extended BBCH scale - general for the descriptions of the growth stages of mono- and dicotyledonous weed species. Weed Research 37: 8 Google Scholar: Author Only Title Only Author and Title

Janssen BJ, Thodey K, Schaffer RJ, Alba R, Balakrishnan L, Bishop R, Bowen JH, Crowhurst RN, Gleave AP, Ledger S, McArtney S, Pichler FB, Snowden KC, Ward S (2008) Global gene expression analysis of apple fruit development from the floral bud to ripe fruit. BMC Plant Biol 8: 16 Google Scholar: Author Only Title Only Author and Title

Jolliffe IT (1986) Principal component analysis. Springer, New York, NY Google Scholar: Author Only Title Only Author and Title

Kudoh H (2016) Molecular phenology in plants: in natura systems biology for the comprehensive understanding of seasonal responses under natural environments. New Phytol 210: 399–412 Google Scholar: Author Only Title Only Author and Title

Labadie M, Denoyes B, Guedon Y (2019) Identifying phenological phases in strawberry using multiple change-point models. J Exp Bot 70: 5687–5701 Google Scholar: Author Only Title Only Author and Title

Leiboff S, Hake S (2019) Reconstructing the Transcriptional Ontogeny of Maize and Sorghum Supports an Inverse Hourglass Model of Inflorescence Development. Current Biology 29: 3410–+ Google Scholar: Author Only Title Only Author and Title

Lorenz DH, Eichhorn KW, Bleiholder H, Klose R, Meier U, Weber E (1995) Growth Stages of the Grapevine: Phenological growth stages of the grapevine (Vitis vinifera L. ssp. vinifera)-Codes and descriptions according to the extended BBCH scale. Grape and wine research 1: 100–103 Google Scholar: Author Only Title Only Author and Title

Massonnet M, Fasoli M, Tornielli GB, Altieri M, Sandri M, Zuccolotto P, Paci P, Gardiman M, Zenoni S, Pezzotti M (2017) Ripening Transcriptomic Program in Red and White Grapevine Varieties Correlates with Berry Skin Anthocyanin Accumulation. Plant Physiol 174: 2376–2396 Google Scholar: Author Only Title Only Author and Title

Menzel A, Sparks TH, Estrella N, Koch E, Aasa A, Ahas R, Alm-KÜBler K, Bissolli P, BraslavskÁ OG, Briede A, Chmielewski FM, Crepinsek Z, Curnel Y, Dahl Å, Defila C, Donnelly A, Filella Y, Jatczak K, MÅGe F, Mestre A, Nordli Ø, PeÑUelas J, Pirinen P, RemiŠOvÁ V, Scheifinger H, Striz M, Susnik A, Van Vliet AJH, Wielgolaski F-E, Zach S, Zust ANA (2006) European phenological response to climate change matches the warming pattern. Global Change Biology 12: 1969–1976 Google Scholar: Author Only Title Only Author and Title

Meyer DH, Schumacher B (2021) BiT age: A transcriptome-based aging clock near the theoretical limit of accuracy. Aging Cell 20 Google Scholar: Author Only Title Only Author and Title

Nagano AJ, Kawagoe T, Sugisaka J, Honjo MN, Iwayama K, Kudoh H (2019) Annual transcriptome dynamics in natural environments reveals plant seasonal adaptation. Nat Plants 5: 74–83 Google Scholar: Author Only Title Only Author and Title

Nagano AJ, Sato Y, Mihara M, Antonio BA, Motoyama R, Itoh H, Nagamura Y, Izawa T (2012) Deciphering and prediction of transcriptome dynamics under fluctuating field conditions. Cell 151: 1358–1369 Google Scholar: Author Only Title Only Author and Title

Nissanka SP, Karunaratne AS, Perera R, Weerakoon WMW, Thorburn PJ, Wallach D (2015) Calibration of the phenology submodel of APSIM-Oryza: Going beyond goodness of fit. Environmental Modelling & Software 70: 128–137 Google Scholar: Author Only Title Only Author and Title

Pagliarani C, Boccacci P, Chitarra W, Cosentino E, Sandri M, Perrone I, Mori A, Cuozzo D, Nerva L, Rossato M, Zuccolotto P, Pezzotti M, Delledonne M, Mannini F, Gribaudo I, Gambino G (2019) Distinct Metabolic Signals Underlie Clone by Environment Interplay in “Nebbiolo” Grapes Over Ripening. Front Plant Sci 10: 1575 Google Scholar: Author Only Title Only Author and Title

Parker AK, De CortÁZar-Atauri IG, Van Leeuwen C, Chuine I (2011) General phenological model to characterise the timing of flowering and veraison of Vitis vinifera L. Australian Journal of Grape and Wine Research 17: 206–216 Google Scholar: Author Only Title Only Author and Title

Parker AK, García de Cortázar-Atauri I, Gény L, Spring J-L, Destrac A, Schultz H, Molitor D, Lacombe T, Graça A, Monamy C, Stoll M, Storchi P, Trought MCT, Hofmann RW, van Leeuwen C (2020) Temperature-based grapevine sugar ripeness modelling for a wide range of Vitis vinifera L. cultivars. Agricultural and Forest Meteorology 285–286 Google Scholar: Author Only Title Only Author and Title

Parker AK, Hofmann RW, van Leeuwen C, McLachlan ARG, Trought MCT (2015) Manipulating the leaf area to fruit mass ratio alters the synchrony of total soluble solids accumulation and titratable acidity of grape berries. Australian Journal of Grape and Wine Research 21: 266–276 Google Scholar: Author Only Title Only Author and Title

Pastore C, Dal Santo S, Zenoni S, Movahed N, Allegro G, Valentini G, Filippetti I, Tornielli GB (2017) Whole Plant Temperature Manipulation Affects Flavonoid Metabolism and the Transcriptome of Grapevine Berries. Front Plant Sci 8: 929 Google Scholar: Author Only Title Only Author and Title

Rabut C (2002) On Pierre Bezier’s life and motivations. Computer-Aided Design 34: 493–510 Google Scholar: Author Only Title Only Author and Title

Robinson S, Glonek G, Koch I, Thomas M, Davies C (2015) Alignment of time course gene expression data and the classification of developmentally driven genes with hidden Markov models. BMC Bioinformatics 16: 196 Google Scholar: Author Only Title Only Author and Title

Sadras VO, Moran MA (2012) Elevated temperature decouples anthocyanins and sugars in berries of Shiraz and Cabernet Franc. Australian Journal of Grape and Wine Research 18: 115–122 Google Scholar: Author Only Title Only Author and Title

Sanchez-Sevilla JF, Vallarino JG, Osorio S, Bombarely A, Pose D, Merchante C, Botella MA, Amaya I, Valpuesta V (2017) Gene expression atlas of fruit ripening and transcriptome assembly from RNA-seq data in octoploid strawberry (Fragaria x ananassa). Sci Rep 7: 13737 Google Scholar: Author Only Title Only Author and Title

Shinozaki Y, Nicolas P, Fernandez-Pozo N, Ma Q, Evanich DJ, Shi Y, Xu Y, Zheng Y, Snyder SI, Martin LBB, Ruiz-May E, Thannhauser TW, Chen K, Domozych DS, Catala C, Fei Z, Mueller LA, Giovannoni JJ, Rose JKC (2018) High-resolution spatiotemporal transcriptome mapping of tomato fruit development and ripening. Nat Commun 9: 364 Google Scholar: Author Only Title Only Author and Title

Theine J, Holtgrawe D, Herzog K, Schwander F, Kicherer A, Hausmann L, Viehover P, Topfer R, Weisshaar B (2021) Transcriptomic analysis of temporal shifts in berry development between two grapevine cultivars of the Pinot family reveals potential genes controlling ripening time. BMC Plant Biol 21: 327 Google Scholar: Author Only Title Only Author and Title

Wang C, Han J, Shangguan L, Yang G, Kayesh E, Zhang Y, Leng X, Fang J (2014) Depiction of Grapevine Phenology by Gene Expression Information and a Test of its Workability in Guiding Fertilization. Plant Molecular Biology Reporter 32: 1070–1084 Google Scholar: Author Only Title Only Author and Title

Wickham H (2009) Ggplot2: Elegant Graphics for Data Analysis, Ed 2nd Edition. Springer Google Scholar: Author Only Title Only Author and Title

